# Transgenerational impact of aberrant inflammation in rat pregnancy

**DOI:** 10.1101/2021.11.09.467505

**Authors:** Takafumi Ushida, Tiziana Cotechini, Nicole Protopappas, Aline Atallah, Charlotte Collyer, Shannyn K. Macdonald-Goodfellow, M. Yat Tse, Louise M. Winn, Stephen C. Pang, Michael A. Adams, Maha Othman, Tomomi Kotani, Hiroaki Kajiyama, Charles H. Graham

**Author notes:** These authors contributed equally to this work (Takafumi Ushida, Tiziana Cotechini).

## Abstract

Children of women with pre-eclampsia have increased risk of cardiovascular (CV) and metabolic disease in adult life. Furthermore, the risk of pregnancy complications is higher in daughters born to women affected by pre-eclampsia than in daughters born after uncomplicated pregnancies. While aberrant inflammation contributes to the pathophysiology of pregnancy complications, including pre-eclampsia, the contribution of maternal inflammation to subsequent risk of CV and metabolic disease as well as pregnancy complications in the offspring remains unclear. Here we demonstrate that 24-week-old female rats (F1) born to dams (F0) exposed to lipopolysaccharide (LPS) during pregnancy (to induce inflammation) exhibited mild systolic dysfunction, increased cardiac growth-related gene expression, abnormal glucose tolerance and coagulopathy; whereas male F1 offspring exhibited abnormal glucose tolerance and increased visceral fat accumulation compared with F1 sex-matched offspring born to saline-treated dams. Both male and female F1 offspring born to LPS-treated dams had evidence of anemia. Fetuses (F2) from F1 females born to LPS-treated dams were growth restricted, and this reduction in fetal growth was associated with increased CD68 positivity and decreased expression of glucose transporter-1 in their utero-placental units. These results indicate that abnormal maternal inflammation can contribute to increased risk of CV and metabolic disease in offspring, and that the effects of inflammation may be transgenerational. This study provides evidence in support of early screening for CV and metabolic disease, as well as pregnancy complications in offspring affected by pre-eclampsia or other pregnancy complications associated with aberrant inflammation.

## Introduction

Pre-eclampsia is a serious condition that affects 3-5% of all pregnancies and is characterized by the development of maternal hypertension and end-organ dysfunction, often accompanied by fetal growth restriction (FGR) after 20 weeks of gestation (1). Moreover, pre-eclampsia is linked to an increased risk of cardiovascular (CV) and metabolic disease in later life for both the affected mothers and their offspring (2–5). There is also evidence that daughters born to women with pre-eclampsia are at increased risk of experiencing pregnancy complications, such as pre-eclampsia, during their reproductive life (6).

According to the Developmental Origins of Health and Disease hypothesis, the fetus undergoes adaptive responses to the hostile *in utero* environment in order to improve survival during complicated pregnancies such as pre-eclampsia and FGR (7–9). While these responses may be beneficial in the *in utero* environment of a complicated pregnancy, they may result in negative consequences following birth, including susceptibility to non-communicable diseases such as hyperlipidemia, hypertension, left ventricular hypertrophy and diabetes later in life (9–11).

There is growing evidence that excessive maternal inflammation during pregnancy is often linked to the etiology of pre-eclampsia (12–14). Many women with pre-eclampsia have a heightened inflammatory state (15–17) and pre-eclampsia is linked to abnormal activation of uterine leukocytes such as macrophages (18, 19), which were also shown to mediate fetal demise in a mouse model (20). Using a model in which pregnant Wistar rats are administered low-dose lipopolysaccharide (LPS) to induce maternal inflammation, we demonstrated that increased levels of tumor necrosis factor alpha are causally linked to the development of features of pre-eclampsia, such as elevated blood pressure, proteinuria, renal abnormalities, impaired spiral artery remodelling, placental hemodynamic alterations, and fetal growth restriction (21). These rats also exhibit long-lasting alterations associated with CV and metabolic disease (22, 23). In the present study, we used the same rat model involving LPS treatment to determine whether aberrant inflammation associated with a pre-eclampsia-like condition also leads to a persistence of CV and metabolic alterations in the offspring. A second objective of our study was to examine whether the effect of aberrant inflammation in pregnancy has transgenerational effects, with negative outcomes in pregnancies from offspring born to LPS-exposed mothers.

## Materials and Methods

### Animal model: F0 and F1 generations

All experiments were conducted in accordance with the guidelines of the Canadian Council on Animal Care. Procedures and protocols were approved by the University Animal Care Committee at Queen’s University. Virgin female and male Wistar rats were purchased from Charles River Laboratories (Montreal, QC, Canada). All animals were provided standard laboratory rat chow and tap water *ad libitum* and were housed at a constant temperature under a 12-h light-dark cycle. Virgin female rats (12-16 weeks of age) were housed with a male rat overnight at a 2:1 ratio, and the presence of spermatozoa in the vaginal smear indicated gestational day (GD) 0.5 for the F0 generation.

Rats were treated as described previously (21, 22). Briefly, F0 pregnant Wistar rats were randomly divided in two groups (N = 6 per group). One group was injected intraperitoneally (i.p.) with LPS (*Escherichia coli* serotype 0111:B4; Sigma-Aldrich Canada, Oakville, ON, Canada) on GD13.5 (10 µg/kg) and daily on GDs 14.5-16.5 (40 µg/kg/day); control rats were injected with saline (1 mL/kg/day) on GDs 13.5-16.5. Rats were allowed to deliver normally, and pups (F1 generation) were culled to eight (four males and four females per litter) on postnatal day (PND) one to control for the potential impact of litter size on final outcomes. Male and female F1 offspring were placed in separate cages at the time of weaning (PND 21). Beginning at 4 weeks of age, weight of male and female F1 offspring was measured weekly ± 1 day. All assessments in F1 offspring were performed at 24 weeks of age unless stated otherwise. The following sections describe procedures performed on the F1 generation, unless specified.

### Mean arterial pressure (MAP) and pulse-wave velocity (PWV) determination

Hemodynamic parameters (blood pressure and PWV) were measured under isoflurane anesthesia and analysed using physiological data analysis software (LabChart version 8, ADInstruments, Colorado Springs, CO, USA) as described previously (22). Aortic PWV is an index of arterial stiffness and is an important parameter of CV disease risk (24, 25). PWV was assessed using the foot-to-foot method described previously (26). Two catheters were inserted into the superior (via left carotid artery) and inferior (via left femoral artery) segments of the aorta using a PE-50 heparinized saline-filled (50 IU/mL) cannula (ID 0.58 mm, OD 0.965 mm; BD Diagnostics, Sparks Glencoe, MD, USA). Both arterial pressure signals were simultaneously monitored and recorded via an attached pressure transducer (Transonic SP200 Pressure System; Transonic Systems Inc., Ithaca, NY, USA). After arterial blood pressure recordings, the distance from the tip of the superior catheter to the inferior catheter was measured. PWV was calculated by dividing the distance between the two catheters over the transmission time. PWV was individually analyzed and averaged using at least 100 normal waveforms. To eliminate the influence of diastolic pressure, β index was calculated according to the following formula: 2.11 x (PWV^2^/Diastolic pressure) (27).

### Echocardiography

Echocardiography was performed to evaluate cardiac structure and systolic and diastolic functions using a Visual Sonics Vevo 2100 Imaging System (Visual Sonics, Toronto, ON, Canada) with a 21-MHz MicroScan transducer (MS-250). Rats were anesthetized with 2-3% isoflurane in air, and heart rate and breathing were monitored. Cardiac function was evaluated in parasternal long axis and apical four-chamber view using M-Mode, colour Doppler Mode and tissue Doppler Mode. Ultrasound data were analyzed using Visual Sonics software (Vevo Lab; Visual Sonics). Tei index (myocardial performance index), which reflects global systolic and diastolic ventricular function, is defined as the sum of isovolumic contraction time and isovolumic relaxation time divided by the ejection time (28).

### Histological assessment of cardiac structure

Following euthanasia, hearts were excised and weighed, and the atria and left ventricles were separated and weighed. The interventricular septum was left attached to the left ventricle. Heart tissues fixed in 10% formalin and paraffin-embedded were sliced into serial 5-µm sections using a Leica RM2125 RTS microtome (Leica Camera AG, Wetzlar, Germany). Sections on slides were stained with hematoxylin and eosin (H&E) to evaluate cardiomyocyte structure, and with Picro-Sirius red (0.1% Direct Red 80 in 1% picric acid, Sigma-Aldrich Canada) to determine collagen content. All slides were analyzed using ImageJ 1.48 software (W.S. Rasband, NIH, Bethesda, MD, USA). In the cross sections stained with H&E, the area of cardiomyocytes in the left ventricles was quantified in each image at 400x magnification. The percent fibrosis area in the left ventricles was quantified in the sections stained with Picro-Sirius red at 100x magnification. Analysis was performed by an observer blinded to the treatments.

### Expression of heart failure- and cardiac growth-related genes

The expression of *Nppa*, (natriuretic peptide type A), *Nppb* (natriuretic peptide type B), *Gata4* (GATA binding protein 4), *Gata6* (GATA binding protein 6), *Ep300* (E1A binding protein p300) and *Mef2c* (myocyte enhancer factor 2C) in left ventricular tissue was assessed by qPCR as reported previously (29). Total RNA was purified from homogenized tissues using a combination of Trizol (Tri Reagent; Molecular Research Centre, Burlington, ON, Canada) and a High Pure RNA isolation kit (Roche Scientific, Laval, QC, Canada) as described previously (30). Generation of cDNA from 1 µg of RNA was performed using a High-Capacity RNA-to-cDNA Kit (Thermo Fisher Scientific, Burlington, ON, Canada). Real-Time PCR was performed using FastStart SYBR Green Master (Roche Scientific) in a LightCycler 480 system II (Roche Scientific). The expression levels of each gene were normalized to β-actin cDNA levels. The sequence of primers for *Nppa, Nppb, Gata4, Gata6, Ep300, Mef2c* and *β-actin* are outlined in Supplementary Table 1. The PCR conditions were as follows: denaturation at 95°C for 5 min, followed by 40-50 cycles at 95°C for 15 s, 62°C for 20 s and 72°C for 20 s.

### Intraperitoneal glucose tolerance test (IPGTT)

IPGTT was performed after overnight fasting to assess ability to metabolize glucose. Rats were injected i.p. with a bolus of 20% glucose solution (2.0 g/kg body weight). Blood glucose levels were measured from tail vein blood using a glucometer (One Touch Ultra 2, Life Scan, Burnaby, BC, Canada) at 0, 15, 30, 60 and 120 minutes after glucose injection.

### Insulin levels and HOMA-IR

Fasting serum insulin (FSI) levels were measured using an ultrasensitive rat insulin ELISA (Mercodia, Uppsala, Sweden) according to the manufacturer’s instructions. Fasting serum glucose (FSG) levels were measured using a glucometer. To determine the level of insulin resistance, the homeostatic model assessment of insulin resistance (HOMA-IR) was calculated according to the following formula: HOMA-IR = [FSG (mmol/L) x FSI (μU/L)]/22.5.

### Complete blood cell (CBC) count and serum analysis

Blood samples were collected at the time of euthanasia. CBC count was performed using an ABC Vet Animal Blood Counter (Scil Animal Care Company, Gurnee, IL, USA) according to the manufacturer’s instructions. Levels of total cholesterol, high-density lipoprotein (HDL) cholesterol, triglycerides (TG), creatinine, and urea were measured in serum samples at the Core Laboratory at Kingston General Hospital. Low-density lipoprotein (LDL) cholesterol was calculated by the formula: total cholesterol - [TG/5 + HDL cholesterol].

### Thromboelastography (TEG)

TEG is a test of whole blood coagulation that provides global information on the dynamics of clot development, stabilization and dissolution (31). Blood coagulation state was evaluated using a TEG 5000 Hemostasis System and TEG Hemostasis Analyser software (Version 4.2) as described previously (32). Briefly, a 340-μL aliquot of citrated whole blood was added to a warm TEG cuvette preloaded with 20 μL of 0.2 mol/L CaCl_2_. TEG parameters including R, K, α angle, MA, CI and LY30 were recorded for approximately 1.0 hour. R is the reaction time until initial fibrin formation, reflecting coagulation factor levels; K is the coagulation time indicating the rapidity of clot formation; α angle is an indication of the rate of clot formation, reflecting fibrinogen activity; maximum amplitude (MA) represents the ultimate strength of the clot, which is an indication of platelet function and fibrinogen activity; coagulation index (CI) is an overall indicator of coagulation; and percent clot lysis at 30 min (LY30) is a measurement of degree of fibrinolysis.

### Induction of pregnancy in the F1 generation and assessment of fetal growth in the F2 generation

Between 10-14 weeks of age, female F1 offspring born to LPS- or saline-treated dams (N = 5 for each group) were mated with non-experimental male rats. Detection of spermatozoa in vaginal lavage indicated GD 0.5. All dams remained untreated until study endpoint on GD 17.5 and fetal wet weights of the F2 generation were assessed at the time of euthanasia.

### Utero-placental assessment of glucose transporter-1 expression

Utero-placental units of F2 offspring (Saline, N = 21; LPS, N = 33) collected from F1 dams (Saline, N = 5; LPS, N = 5) at GD 17.5 were fixed using 4% paraformaldehyde, processed, embedded in paraffin, and sectioned. Tissue sections (5-μm) were rehydrated using a graded ethanol series and subjected to heat-mediated citrate antigen retrieval (BioGenex, Freemont, CA, USA) for 12 minutes. Endogenous tissue peroxidases were blocked by immersion in a 0.6% hydrogen peroxide solution in methanol for 20 minutes. Following protein blocking (1% normal goat serum for 30 minutes at room temperature), anti-rabbit glucose transporter-1 (Glut-1; Invitrogen; PA5-32428, 1:200) was added to tissues and incubated for two hours at room temperature. Detection of the primary antibody was achieved by incubation of tissues with a species-specific secondary polymer (Histofine^®^ Simple Stain MAX PO; Nichirei Biosciences Inc., Tokyo, Japan) for 30 minutes at room temperature, followed by incubation with 3,3’-diaminobendizine (DAB; Cell Signaling Technology; 8059S). Sections were counterstained using Gill’s hematoxylin (Thermo Fisher Scientific), dehydrated, mounted, and scanned using an Olympus VS120 high resolution scanner. Digital image analysis was performed using HALO^®^ image analysis platform (Indica Labs). The labyrinth, junctional zone and mesometrial triangle of utero-placental units were manually annotated and the percent strong positive staining was calculated for each region.

### Immunohistochemical detection and analysis of utero-placental CD68 Expression

F2 utero-placental units (Saline, N = 18; LPS, N = 21) collected from F1 dams (Saline, N = 5; LPS, N = 5) at GD 17.5 were fixed using 4% paraformaldehyde, processed, embedded in paraffin, and sectioned. Tissue sections (5-μm) were subjected to immunohistochemistry for CD68 expression and analyzed using a similar approach as described for glucose transporter-1, with the following modifications: protein blocking with goat serum was carried out for 45 minutes and sections were incubated overnight in a humidified chamber at 4°C with anti-CD68 primary antibody (ab125212, Abcam; 1:350). Digital analysis was performed using Halo® image analysis platform (Indica Labs, Albuquerque, NM, USA). Scanned sections of utero-placental units were manually annotated into the mesometrial triangle, labyrinth, and junctional zone by a blinded observer. The percent strong and moderate positive staining was calculated for each region.

### Polychromatic flow cytometry assessment of bone marrow from F1 offspring

Bone marrow flushed from femurs from male and female F1 offspring at 12 and 24 weeks of age was stained using two different cocktails of antibodies (Supplementary Table 2) to evaluate the lymphoid and myeloid immune compartments. Briefly, cells (1 x 10^6^) were plated in buffer (PBS with 0.5% bovine serum albumin and 1 mM EDTA) and non-specific Fc binding was blocked via incubation with mouse anti-rat CD32 (BD Biosciences #550271; 1:200) for 30 minutes on ice. Cells were then stained with 100 μL of primary antibody cocktail for 30 minutes on ice, washed, and fixed for 12 minutes (BD Biosciences #554655). Data were acquired on a CytoFLEX S flow cytometer (Beckman Coulter) and analyzed using FlowJo version 10.7 software (BD Biosciences). Cells of myeloid (Supplementary Fig. 1*a*) and lymphoid (Supplementary Fig. 1*b*) lineages were assessed between treatment groups over the two time-points.

### Statistical Analysis

Statistical analysis was performed using GraphPad Prism 9.0 software (GraphPad Software Inc., La Jolla, CA, USA). When comparing two groups, Student’s t test and Mann-Whitney U test were performed for parametric and non-parametric data, respectively. Repeated measures two-way ANOVA with Sidak’s multiple comparisons test was used to statistically analyze data on body weight, cardiac parameters, food intake, glucose levels following IPGTT, and TEG parameters in male and female F1 offspring born to LPS and saline treated F0 dams. Fisher’s Exact test was used to compare the proportion of FGR offspring. Two-way ANOVA followed by Tukey’s *post hoc* analysis was performed to assess differences in bone marrow immune cell composition in F1 male and female rats at 12 and 24 weeks of age. Rout’s test was performed to remove outliers. Data are expressed as means ± SEM. A value of *p* < 0.05 was considered statistically significant.

## Results

### Effect of aberrant maternal inflammation on neonatal outcomes

To assess whether aberrant maternal inflammation affects neonatal outcomes, we defined FGR in our rats as a birth weight falling below the 10^th^ percentile (33). Aberrant inflammation resulted in a significant increase in the FGR rate of the female F1 generation and a significant reduction in birth weight of male and female F1 offspring born to LPS-treated F0 dams (Table 1). There was no significant difference in mortality rates for offspring born to either saline or LPS-treated mothers during the neonatal period (data not shown).

### Effect of aberrant maternal inflammation on offspring cardiac function and structure

MAP was comparable for the two groups of both male and female F1 offspring (Fig. 1*a*). As adults, F1 female offspring born to LPS-treated dams exhibited a trend towards decreased ejection fraction (EF; Fig. 1*b*; *p* = 0.06) and fractional shortening (FS; Fig. 1*c*; *p* = 0.057) compared with adult F1 female offspring born to control mothers. No differences in global systolic and diastolic function (Tei index; Supplementary Fig. 2*a*), pulse-wave velocity (Supplementary Fig. 2*b*) or β index (Supplementary Fig. 2*c*) were observed between groups. Details of all cardiac parameters are shown in Supplementary Table 3.

**Figure 1.**
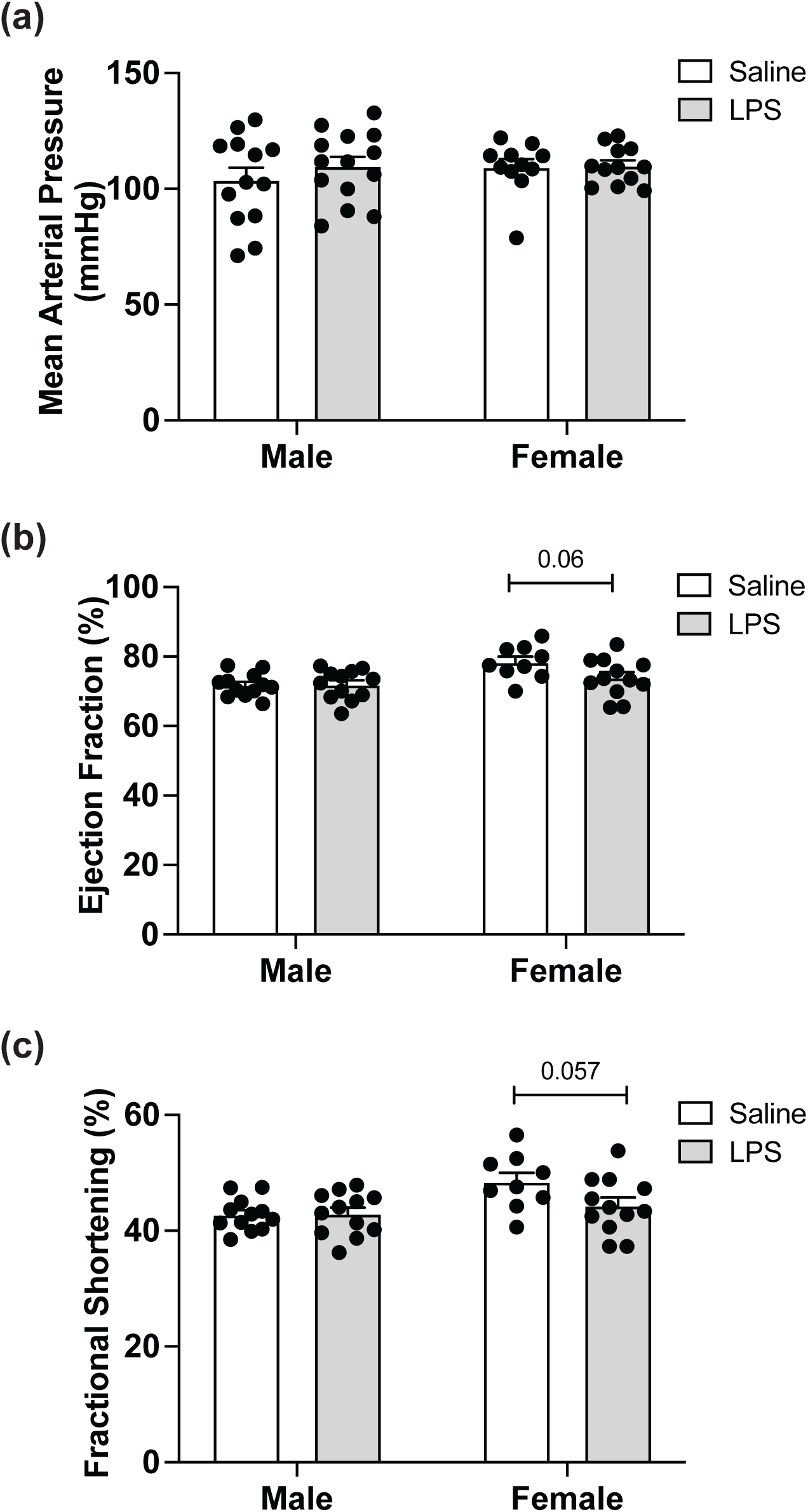
*Effect of aberrant maternal inflammation on F1 offspring cardiac function and structure.* Mean arterial pressure *(a)* was measured by inserting two catheters into the left carotid artery and left femoral artery whereas ejection fraction *(b)*, and fractional shortening *(c)* were assessed by echocardiography. MAP, mean arterial pressure. Data are presented as mean ± SEM. Each filled circle represents an individual rat.

To gain further insight into the mild systolic dysfunction observed in F1 female offspring at 24 weeks of age, we measured heart weights and calculated various ratios, assessed cardiac histology, and expression of genes related to cardiac development. F1 female offspring born to LPS-treated dams exhibited significantly increased cardiac mRNA levels of myocyte enhancer factor-2c (Mef2c) measured at 24 weeks of age (Fig. 2*a*). No significant differences were found in the expression levels of *Nppa, Nppb, Gata4, Gata6,* and *Ep300* (Fig. 2*b-f*). Similarly, no significant differences were found in cardiomyocyte area (Supplementary Fig. 3*a*) or width (Supplementary Fig. 3*b*), and interstitial fibrosis area (Supplementary Fig. 3*c*) in offspring of control versus LPS-treated dams. Finally, no significant differences were observed between treatment groups when comparing heart weight-to-tibia length ratios, heart weight-to-body weight ratios, or right ventricle to left ventricle plus septum weight ratios (data not shown).

**Figure 2.**
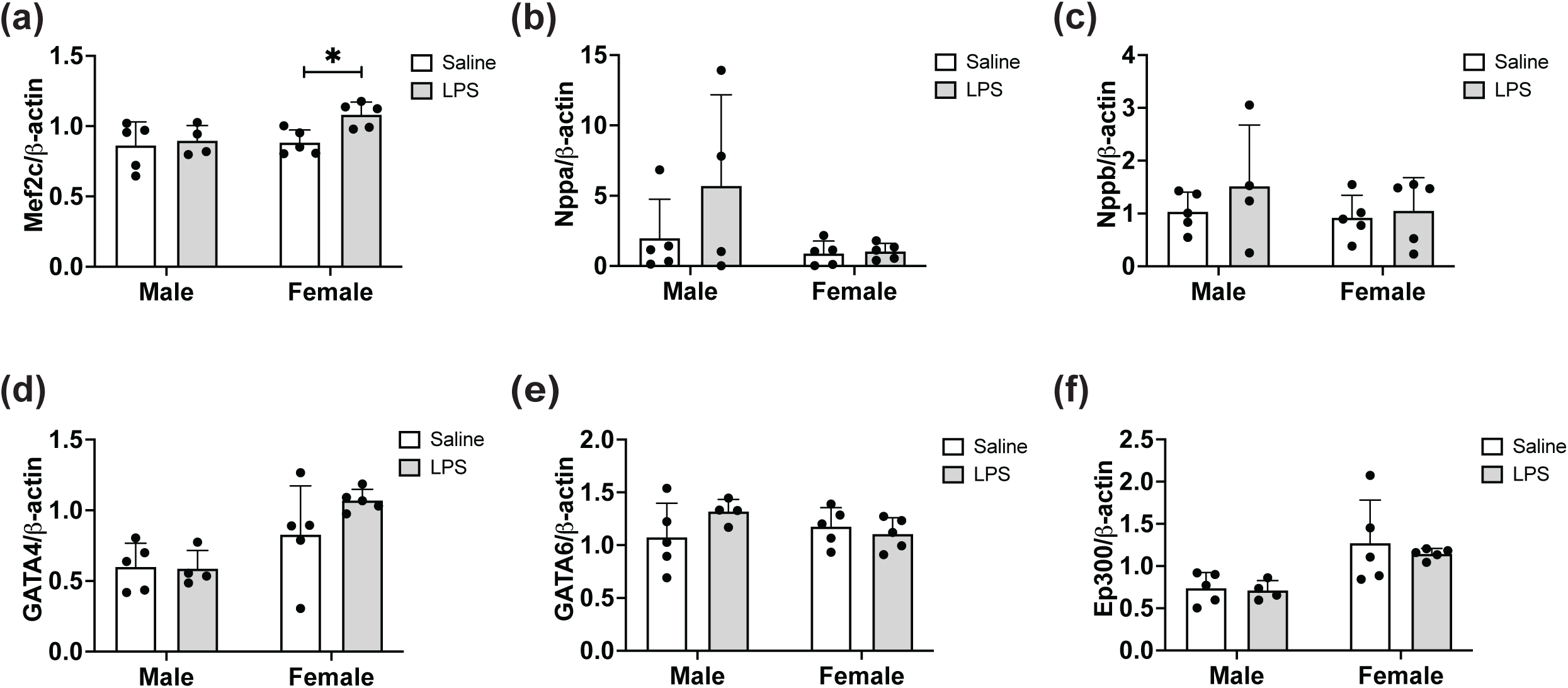
*Effect of aberrant maternal inflammation on F1 offspring heart failure- and cardiac growth-related gene expression.* Left ventricular mRNA expression of *Mef2c (a)*, *Nppa (b)*, *Nppb (c)*, *Gata4 (d)*, *Gata6 (e)* and *Ep300 (f)* were measured by qPCR. Data are presented as mean ± SEM; \**p* < 0.05. Each filled circle represents an individual rat.

### Effect of aberrant maternal inflammation on offspring metabolic functions

Although the post-weaning weights of male F1 pups from LPS-treated dams was not significantly different from weights of F1 pups born to control-treated mothers, two-way ANOVA analysis revealed that LPS treatment significantly reduced post-weaning weights of F1 female pups in the LPS treatment group compared with weights of F1 offspring from saline-treated controls (Fig. 3*a*). This difference was not attributable to differences in food intake as there were no observed differences in food consumption between male or female F1 offspring from born to either saline or LPS-treated mothers (Fig. 3*b*). To determine the effect of maternal inflammation on glucose tolerance in adult F1 offspring, IPGTT was performed at 24 weeks of age. Compared with F1 controls, both F1 adult male and female offspring born to LPS-treated mothers exhibited a significant increase in glucose levels 15 minutes following i.p. bolus glucose injection (Fig. 3*c, d*). Neither fasting insulin levels (Fig. 3*e*) nor insulin resistance (HOMA-IR; Fig. 3*f*) measured at 24 weeks of age were significantly different in F1 offspring of saline vs. LPS-treated dams.

**Figure 3.**
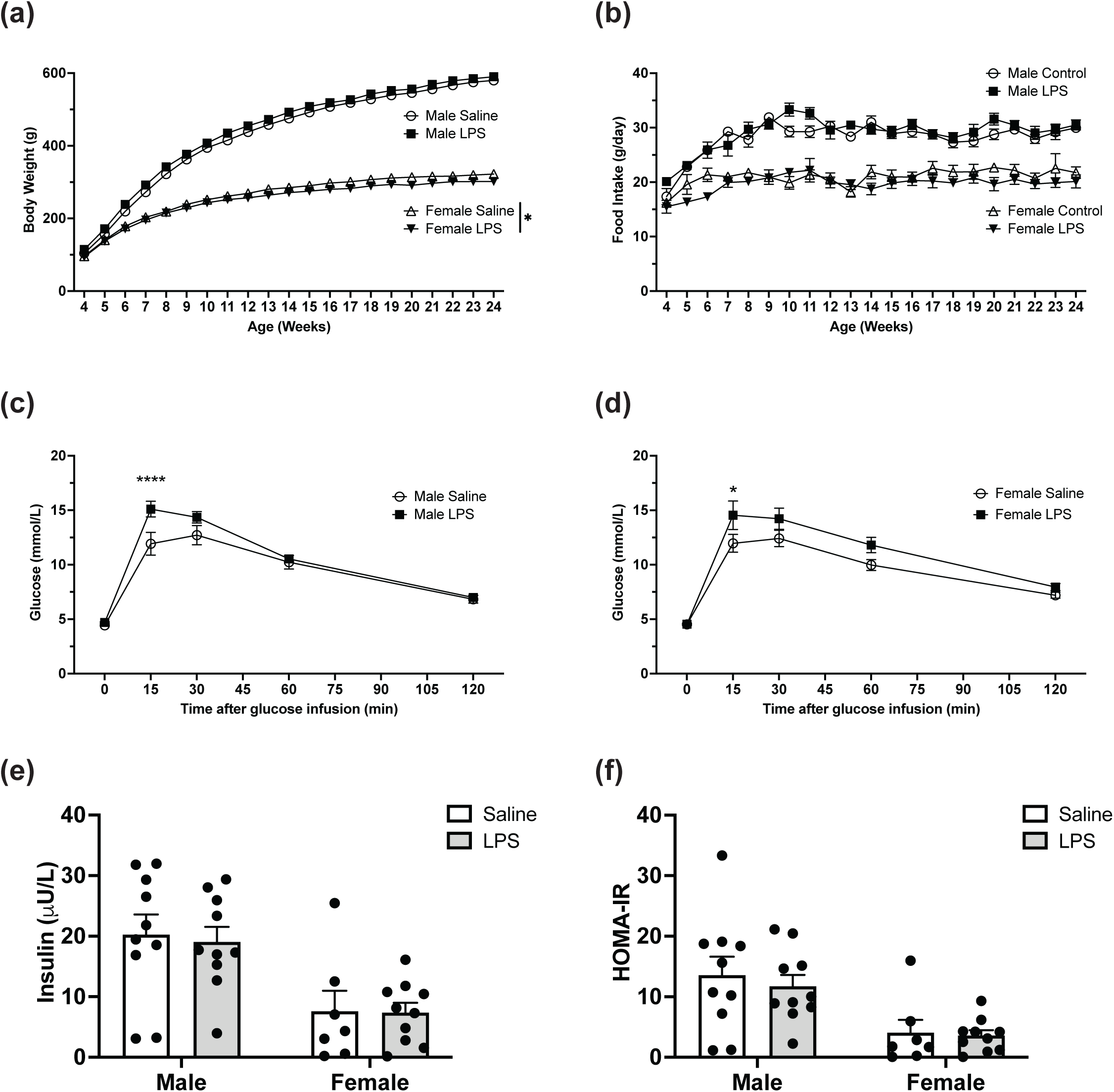
*Effect of aberrant maternal inflammation on F1 offspring metabolic functions.* Body weight in male and female offspring (*a*; n = 13-17) and food consumption *(b)* was measured weekly ± 1 day (n = 12-17). Intraperitoneal glucose tolerance test was performed at 24 weeks of age *(c and d)*. Blood glucose levels were assessed after injection of 20% glucose solution (injected at time 0) (n = 11-15). Fasting insulin level was measured at 24 weeks of age (*e*; n = 10). HOMA-IR was assessed at 24 weeks of age (*f*; n= 10). Data are presented as mean ± SEM; \**p* < 0.05; \*\*\*\**p* < 0.001.

### Effect of aberrant maternal inflammation on offspring fat mass and lipid profile

Visceral fat accumulation was measured at 24 weeks of age. In adult male F1 offspring from LPS-treated dams, omental fat accumulation trended toward being significantly increased compared with saline controls (Fig. 4*a*; *p* = 0.077). Mesenteric (Fig. 4*b*, epididymal (Fig. 4*c*) and retroperitoneal fat (Fig. 4*d*) masses were significantly increased in male F1 offspring from LPS-treated dams. In contrast, no differences were observed in visceral fat accumulation for adult female F1 offspring between groups (Fig. 4*a-d*). No significant difference was found in the circulating lipid profile at 24 weeks of age in F1 male or female offspring born to saline or LPS-treated dams (Fig. 4*e*).

**Figure 4.**
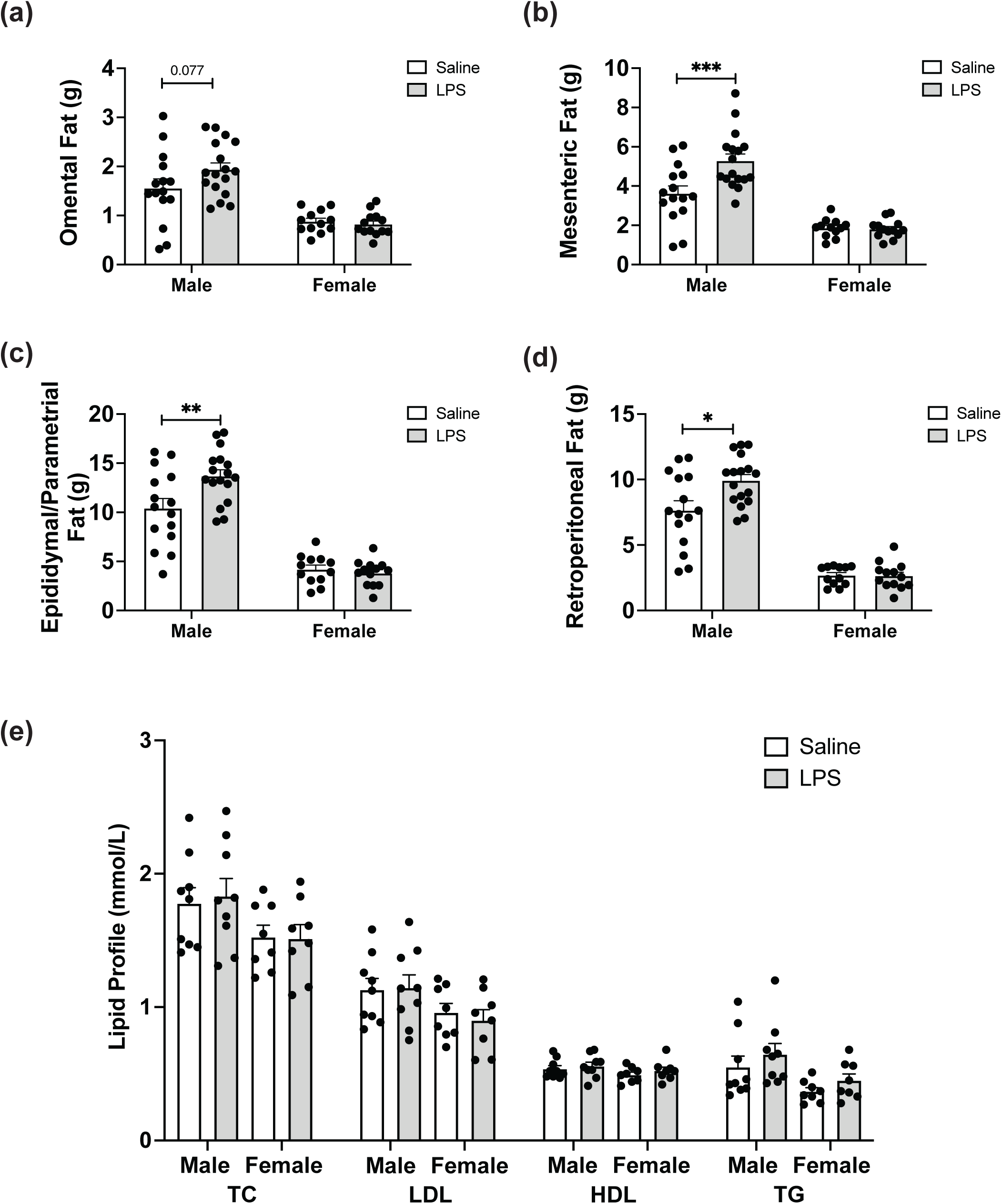
*Effect of aberrant maternal inflammation on F1 offspring fat mass and lipid profile.* Visceral fat accumulation was measured at 24 weeks of age including omental fat *(a)*, mesenteric fat *(b)*, epididymal/parametrial fat *(c)* and retroperitoneal fat *(d)*. Serum lipids (total cholesterol, low density lipoprotein, high density lipoprotein and triglycerides) were assessed at 24 weeks of age *(e)*. Data are presented as mean ± SEM; \**p* < 0.05; \*\**p* < 0.01; \*\*\**p* < 0.001. Each filled circle represents an individual rat. TC, total cholesterol; LDL, low-density lipoprotein; HDL, high-density lipoprotein; TG, triglycerides.

### Effect of aberrant maternal inflammation on offspring complete blood count and renal function

At 24 weeks of age, no differences in circulating leukocyte numbers were observed (Fig. 5*a*) in F1 offspring. However, male F1 offspring born to LPS-treated dams exhibited a trend toward significantly decreased red blood cell (RBC) numbers (Fig. 5*b*; *p* = 0.085) and hematocrit (Fig 5*c*; *p* = 0.07); whereas, compared with adult female rats born to saline-treated dams, female F1 offspring born to LPS-treated mothers had significantly decreased RBC numbers (Fig. 5*b*) and hematocrit (Fig 5*c*), and showed a trend toward a reduction in hemoglobin levels (Fig 5*d*; *p* = 0.056). To investigate the cause of anemia, mean corpuscular volume (MCV) and mean corpuscular hemoglobin (MCH) were calculated and were found to be comparable in both male and female F1 offspring born to either control or LPS-treated rats (data not shown), thus indicating normochromic normocytic anemia. Kidneys collected from adult male F1 offspring born to LPS-treated mothers exhibited a trend toward weighing less than kidneys collected from adult male F1 offspring of saline-treated controls (Fig. 5*e*; *p* = 0.057). No significant differences in renal function as measured by serum creatinine (Fig. 5*f*) or urea (Fig. 5*g*) were observed.

**Figure 5.**
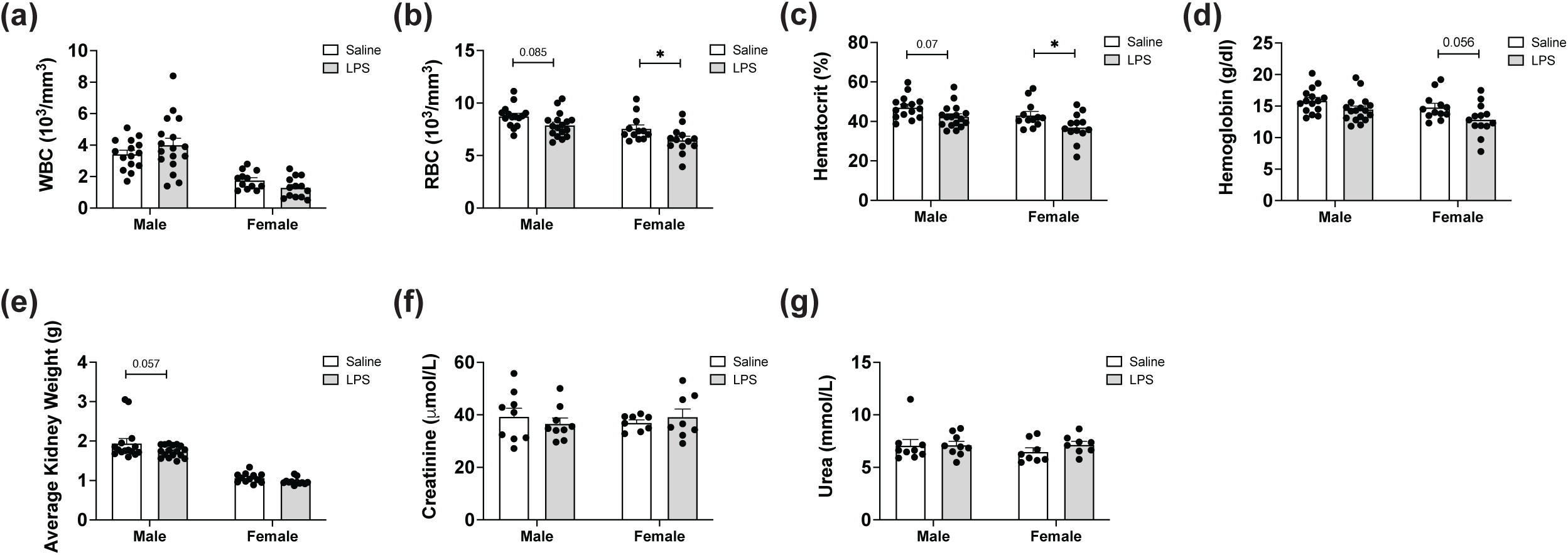
*Effect of aberrant maternal inflammation on complete blood counts and renal function of F1 offspring.* White blood cell counts *(a)* red blood cell counts *(b)*, hematocrit *(c)* and hemoglobin levels *(d)* assessed from whole blood at 24 weeks of age. Average kidney weights *(e)* and serum creatinine *(f)* and urea *(g)* were measured at 24 weeks of age. Data are presented as mean ± SEM; \**p* < 0.05. Each filled circle represents an individual rat. WBC, white blood cell; RBC, red blood cell.

### Effect of aberrant maternal inflammation on offspring coagulation function

Thromboelastography (TEG) was performed on citrated, whole blood collected at the time of euthanasia (Table 2). Two-way ANOVA revealed that LPS treatment and not sex significantly affected reaction time [R], clot formation time [K] and rate of clot formation [α angle]. Specifically, compared with the adult female F1 offspring from saline-treated dams, adult F1 female offspring of LPS-treated dams showed significantly increased clot formation time (K).

### Transgenerational effect of aberrant maternal inflammation on fetal growth restriction

The average weight of F2 fetuses from F1 adult female offspring born to LPS-treated dams (0.717g ± 0.013g) was significantly decreased compared with the average weight of F2 fetuses from F1 adult female offspring born to saline-treated dams (0.770g ± 0.010g; Table 3). The proportion of growth restricted F2 fetuses from adult rats born to F1 LPS-treated dams (31.7%) was significantly increased compared with the proportion of FGR F2 fetuses from adult F1 rats born to saline-treated dams (10%; Table 3).

### Assessment of Glut1 and CD68 expression in F2 utero-placental units

Glut1 expression was significantly reduced in the mesometrial triangle (Fig. 6) and labyrinthine zone of F2 utero-placental units collected from F1 dams born to LPS-treated F0 dams compared with saline controls. No difference was observed in the density of Glut1 expression within the junctional zone. CD68 positivity was significantly increased in the mesometrial triangle, junctional zone and labyrinth regions of F2 utero-placental units from to LPS-treated F0 dams compared with saline controls (Fig. 7). This observed increase in the number of CD68^+^ macrophages was not associated with changes in the bone marrow myeloid compartment from F1 offspring (Supplementary Table 4).

**Figure 6.**
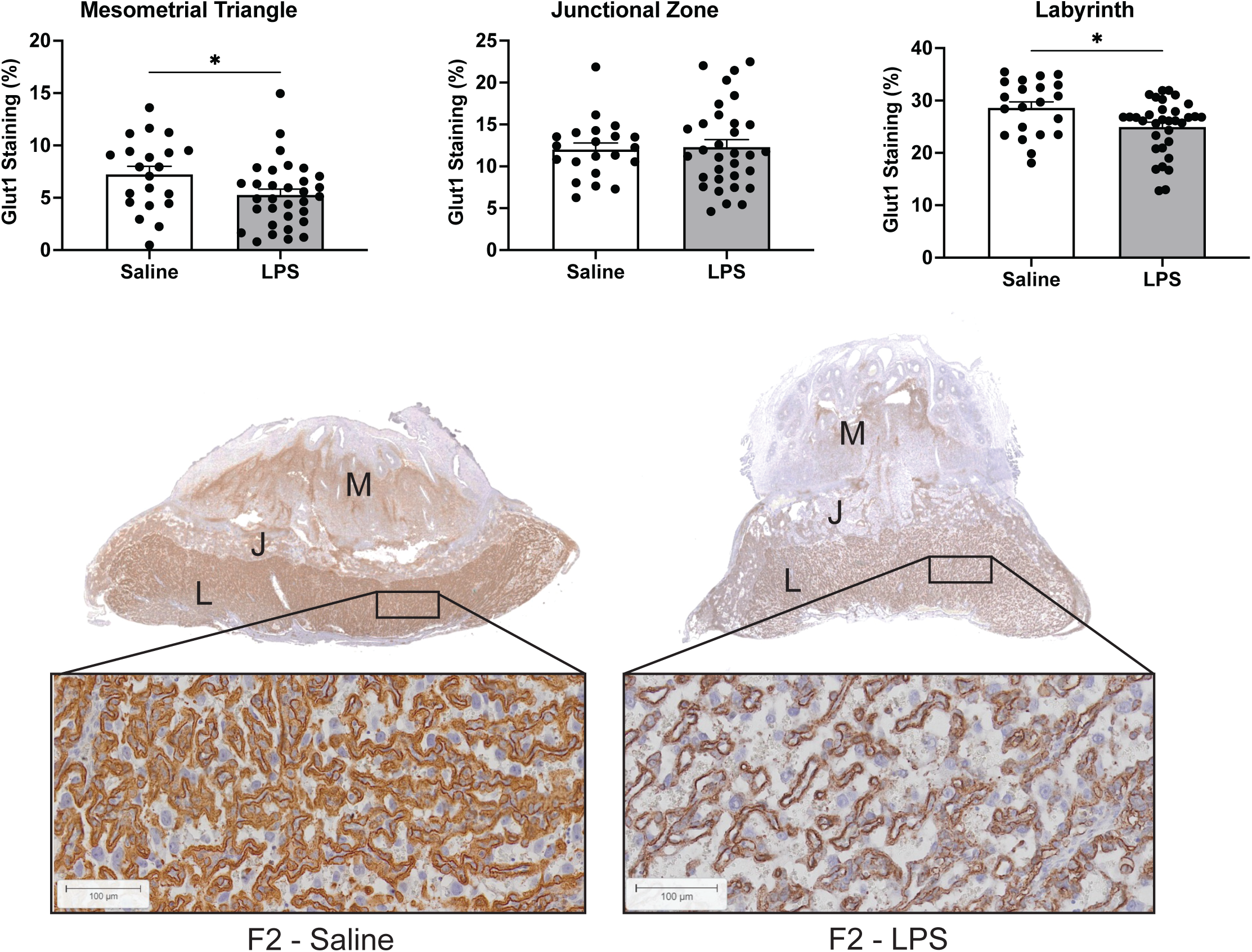
*Glucose transporter-1 expression in utero-placental units of the F2 generation.* Immunolocalization of GLUT-1 expression in the mesometrial triangle, junctional zone and labyrinth of F2 utero-placental units collected from F1 pregnant rats (N=5/group) born to either saline- or LPS-treated F0 dams. Data are presented as mean ± SEM; \**p* < 0.05. Each filled circle represents an individual utero-placental unit. M, mesometrial triangle; J, junctional zone; L, labyrinth.

**Figure 7.**
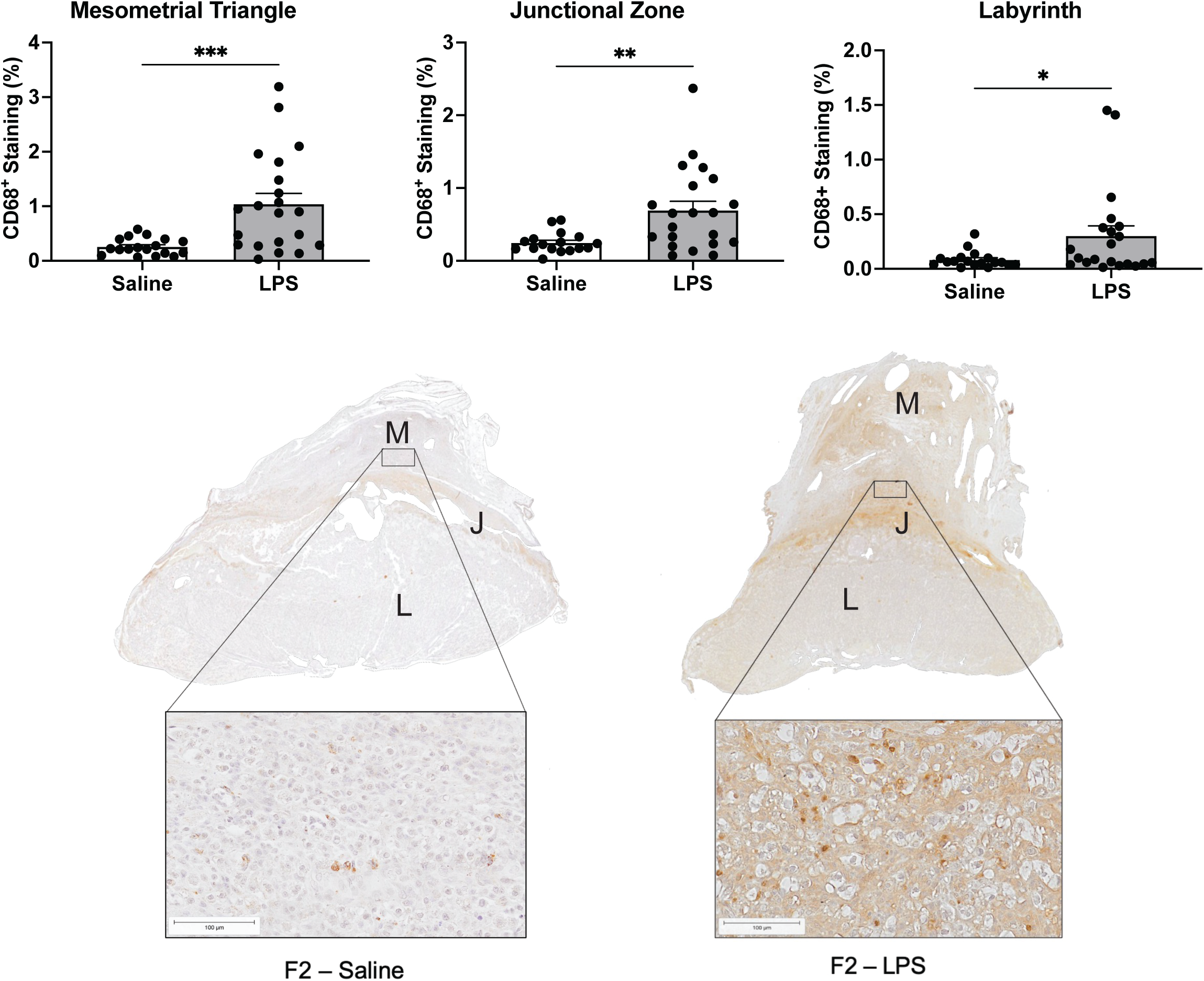
*CD68 expression in utero-placental units from the F2 generation.* Immunolocalization of CD68 expression in the mesometrial triangle, junctional zone and labyrinth of F2 utero-placental units collected from F1 pregnant rats (N=5/group) born to either saline- or LPS-treated F0 dams. Data are presented as mean ± SEM; \**p*<0.05. Each filled circle represents an individual utero-placental unit. M, mesometrial triangle; J, junctional zone; L, labyrinth.

## Discussion

This study revealed that abnormal inflammation in pregnancy results in the acquisition of long-lasting cardiovascular and metabolic disease risk factors in F1 offspring and impairs *in utero* growth of both the F1 and F2 generations. Overall, our results provide evidence in support of the concept that excessive inflammation is a key factor contributing to the increased risk of disease in children of women afflicted by complications such as pre-eclampsia, and that the effect of aberrant inflammation in pregnancy may have transgenerational consequences.

Several animal studies have demonstrated sex-specific responses to prenatal adverse conditions (34). Most studies reveal that male offspring are more sensitive than females to adverse intrauterine conditions and therefore more prone to adult-onset disease (34). Our study provides evidence in support of sex-specific responses to *in utero* exposure to maternal inflammation as only F1 male offspring of LPS-treated rats exhibited higher fat accumulation compared with controls. In contrast, mild systolic dysfunction was only observed in F1 female offspring of LPS-treated rats, which is consistent with a previous study showing systolic dysfunction in response to cardiac damage induced by isoproterenol in female versus male rats (35).

In our study, LPS administration to pregnant dams resulted in significantly reduced *in utero* growth for both male and female offspring. Whereas male offspring from LPS- treated dams exhibited catch-up growth, female offspring from LPS-treated dams had significantly lower body weight between 4 and 24 weeks of age compared with controls. In a rat model of pre-eclampsia induced after reducing the uterine perfusion pressure (RUPP model), male and female offspring exhibited growth restriction and did not undergo catch-up growth (36). Similarly, there was no catch-up growth in male mouse offspring five months following a pre-eclampsia-like condition induced after inoculation with an adenovirus that expresses soluble fms-like tyrosine kinase-1 (sFlt-1); female offspring, however, showed catch-up growth at two months of age (37). In contrast, in a rat model of FGR induced by under-nutrition during pregnancy, both male and female offspring showed catch-up growth by 12-30 weeks of age (38). In our inflammation-induced model of pre-eclampsia, rats exhibited reduced maternal body weight gain during pregnancy, indicating similarities with the rat under-nutrition model of FGR. Thus, it is plausible that the mismatch in prenatal and post-natal nutrition accelerates catch-up growth during lactation in our model; however, further investigation is required to determine the reasons for the sex-specific difference in growth rate.

Our results differ from previous human and animal studies of pre-eclampsia in that the offspring from our LPS-treated rats did not exhibit clear evidence of hypertension and cardiac hypertrophy. A systematic review reported that children of women with pre-eclampsia had elevated systolic blood pressure during childhood and early adulthood (2). Using the sFlt-1 over-expression mouse model of pre-eclampsia, Lu *et al.* showed elevated MAP only in male offspring at six months of age (37). Similarly, only the male offspring of RUPP dams exhibited a significant increase in MAP at 12 weeks of age (36). It is remarkable that the increase in MAP in that model was substantially higher than what has been observed in humans. In our present study, both male and female offspring of LPS-treated dams had lower MAP at 24 weeks of age compared with the previous observations in animal studies. In those previous two animal studies, blood pressure was measured by implantable radiotelemetry in conscious animals, resulting in continuous and precise MAP recordings (36, 37). In contrast, we measured blood pressure in rats at a single time point prior to euthanasia using a carotid artery catheter inserted under isoflurane anesthesia, which has known blood pressure-lowering effects (39). The sensitivity of this approach may not detect small changes in MAP, which could explain why we did not see differences between the two groups. It is also possible that in an inflammation-based rat model of pregnancy complications, such as ours, hypertension in the offspring develops after 24 weeks of age.

Using echocardiography, we found a trend toward mild systolic dysfunction in female offspring from LPS-treated rats. Although only a few studies have focused on the long-term effect of maternal inflammation on offspring cardiac function, our finding is consistent with a previous report that showed left ventricular systolic dysfunction in mice at eight weeks of age following maternal administration of LPS (80 μg/kg i.p) on GD 16 (40). In contrast, another study showed increased systolic function in 20-week-old Sprague Dawley rats born to dams treated with LPS (0.79 mg/kg) on GD 8, 10 and 12 (35). Another similar study revealed diastolic dysfunction in Sprague Dawley rat at four months of age after maternal LPS (0.79 mg/kg) administration on GD 8, 10 and 12 (41). It is well recognized that the sensitivity to LPS varies according to the model, with Wistar rats being more sensitive to LPS compared to Sprague Dawley rats (42). Thus, it is possible that the observed discrepancies on the reported effects of maternal inflammation on offspring cardiac function are due to rat strain-specific responses, as well as differences in the timing and dose of LPS administration.

A previous study demonstrated that systemic maternal inflammation and neonatal hypoxia induces cardiac remodelling and left ventricular dysfunction in mice (40). Other animal studies have shown that prenatal chronic hypoxia can cause cardiac dysfunction in the offspring as well as increased susceptibility to ischemia-reperfusion injury in the heart during adult life (43, 44). Our rat model of LPS-induced inflammation demonstrated increased spiral artery resistance index and placental oxidative stress, indicative of impaired placental perfusion, on GD 17.5 (21, 45, 46). Under such conditions, prenatal hypoxia during the second half of pregnancy could lead to cardiac remodelling in the offspring. As we expected, female offspring at 24 weeks of age showed increased expression of *Mef2c*, a transcription factor associated with cardiac hypertrophy and remodelling (29, 47).

F1 offspring of LPS-treated dams at 24 weeks of age had evidence of normochromic normocytic anemia, a risk factor for CV disease, and in particular chronic heart failure (48). This type of anemia is not caused by under-nutrition such as iron deficiency and vitamin insufficiency, which often lead to microcytic hypochromic anemia and macrocytic anemia, respectively (49). Although we suspected renal anemia manifested as normochromic normocytic anemia (49), due to the reduced kidney size in offspring from LPS-treated dams, we could not find any renal dysfunction after analysis of serum samples. Further studies are required to fully elucidate the cause of anemia in these rats; however, it is possible that anemia, together with the increased expression of cardiac growth-related genes, contributed to the development of the mild systolic dysfunction observed in the female offspring.

Our study also revealed abnormal glucose tolerance in both male and female offspring from LPS-treated rats at 24 weeks of age; however, there were no significant differences in the lipid profile. Interestingly, our results are consistent with those obtained using sFlt-1 overexpressing mice (50). Female offspring from sFlt-1 overexpressing mice displayed a higher glucose response to IPGTT compared with controls, and male offspring showed lower fasting insulin level at six months of age, suggesting abnormal glucose metabolism (50). According to a systematic review and meta-analysis, no significant difference was observed in lipid profile in umbilical cord blood or infant blood collected following pre-eclamptic versus uncomplicated pregnancies (5).

Our present study revealed an effect of LPS treatment on TEG parameters associated with prolonged clot formation measured in F1 offspring at 24 weeks of age. In humans, coagulopathy is strongly linked to the development of CV disease, especially a hypercoagulable status with suppression of fibrinolysis (51). However, our TEG results indicated a hypocoagulable state. We have previously used TEG with success to investigate coagulopathies in various studies using a similar rat model (32, 45, 46, 52).

The hypocoagulable state observed in the present study may be a consequence of endothelial dysfunction associated with the mild systolic dysfunction and metabolic alterations observed in the F1 offspring, which can impair the synthesis and/or release of both procoagulant and anticoagulant factors (53, 54). Anemia is another potential explanation for the hypocoagulable state; however, only a few studies have analyzed the effect of anemia on coagulation parameters (55, 56). Brooks *et al*. revealed that low hematocrit leads to high K and low R and α angle (55). In contrast, Roeloffzen *et al*. showed anemia leads to increased R and α angle (56). Thus, the effect of anemia on coagulation and whether there is a mechanistic link between hypocoagulability and CV disease remain unclear.

In our study, we also observed a transgenerational impact of inflammation-induced impaired fetal growth. These observations align with human data demonstrating that daughters born to mothers with pre-eclampsia experience an increased risk of developing pregnancy complications (57). Moreover, using a rat model of utero-placental insufficiency, Gallo and colleagues reported impaired second-generation fetal growth when their mothers were born small for gestational age (58). Our rat model of inflammation-induced fetal growth restriction was previously shown to be associated with reduced spiral artery remodelling, utero-placental insufficiency and increased CD68^+^ immunoreactivity in the mesometrial triangle (21). In the present study, we report increased CD68^+^ positivity in F2 utero-placental units from fetuses who are descendants of LPS-treated grandmothers compared to controls. This is notable given that female F1 offspring from LPS-treated rats were not directly exposed to LPS during their pregnancies with the F2 generation, and yet CD68^+^ immunoreactivity was increased across the utero-placental units of the F2 generation. This increased presence of CD68^+^ cells was not attributable to alterations in the myeloid compartment of bone marrow of F1 offspring from LPS-treated dams. It is possible, therefore, that this observed increase in the numbers of CD68^+^ cells is due to proliferation of resident myeloid populations. Presence of large numbers of CD68^+^ cells is suggestive of a pro-inflammatory state, which in turn is linked to altered placental nutrient transport (59). To our knowledge, the current work is the first to describe a transgenerational link between inflammation-induced pregnancy complications in the F0 generation and reduced growth of the F2 generation.

The developing neonate relies on sufficient glucose uptake from maternal blood to achieve its genetically pre-determined growth potential. The glucose transporter GLUT1 is expressed by various trophoblast lineages in both human and rodent pregnancy and is the primary transporter responsible for glucose transfer across the placenta (60–62). Our findings that GLUT1 expression is reduced in the labyrinth zone in utero-placental units of second-generation descendants of LPS-treated dams is consistent with other work reporting a link between reduced labyrinth GLUT1 expression and impaired fetal growth (63, 64). Moreover, a recent human study revealed that maternal obesity, a condition associated with chronic low-level inflammation, was linked to decreased placental GLUT1 expression (65). Though the results of our study do not identify a causal mechanism through which FGR is heritable across generations, our data support an association between inflammation-induced FGR and reduced GLUT-1 expression in the placenta of subsequent generations.

In conclusion, this study provides novel evidence that aberrant maternal inflammation associated with fetal growth restriction leads to persistent physiological alterations in the F1 generation, some of which are known to increase the risk of acquiring cardiovascular and metabolic disease in adult life. We also show that adult female F1 offspring born to LPS-treated dams experience pregnancy complicated by FGR. Thus, pregnancy may act as a ‘second hit’ that when superimposed upon the above-mentioned latent phenotypes in the F1 generation, results in growth restriction of the F2 generation. In a previous study using the same animal model we demonstrated that aberrant inflammation in pregnancy also leads to persistence of risk factors for cardiovascular and metabolic disease in the affected mothers (22). Therefore, our findings support the concept that developing strategies to prevent or manage pregnancy complications associated with abnormal inflammation may significantly reduce the burden of subsequent disease in mothers and subsequent generations.

## Acknowledgements

We would like to thank Kim Laverty for technical support with the assessments of PWV and blood pressure; Rebecca Maciver for assistance with echocardiography; Terry Yantian Li for technical support with histology and PCR; Dr. Lori Minassian for her technical expertise and assistance with PCR; Allegra Quadri for blinded analysis of histology; Deborah Harrington and Dr. Janine Handforth for determining complete blood counts; Spencer Barr for technical support with western blotting and TEG; and Maggie Chasmar for assistance with animal care.

## Financial Support

This work was supported by grants from the Canadian Institutes of Health Research (CIHR: MOP-119496 and PJT-156222) awarded to C.H. Graham and from the Japan Society for the Promotion of Science (JSPS KAKENHI 19K18637) awarded to Takafumi Ushida.

## Conflict of interest

The authors have no conflicts of interest to declare in association with this study.

## Author Contributions

Study conception/design: TU, TC, MAA, LMW, CHG,

Primary data acquisition: TU, TC, NP, AA, CC, SKM-G, MYT, SCP

Data analysis and interpretation: TU, TC, NP, AA, CC, SKM-G, MYT, MO, MAA, TK, HK, CHG

Manuscript draft: TU, TC, AA, CC, CHG

Manuscript revisions and final editing: All authors

## Supplementary Figures

**Supplementary Figure 1.**
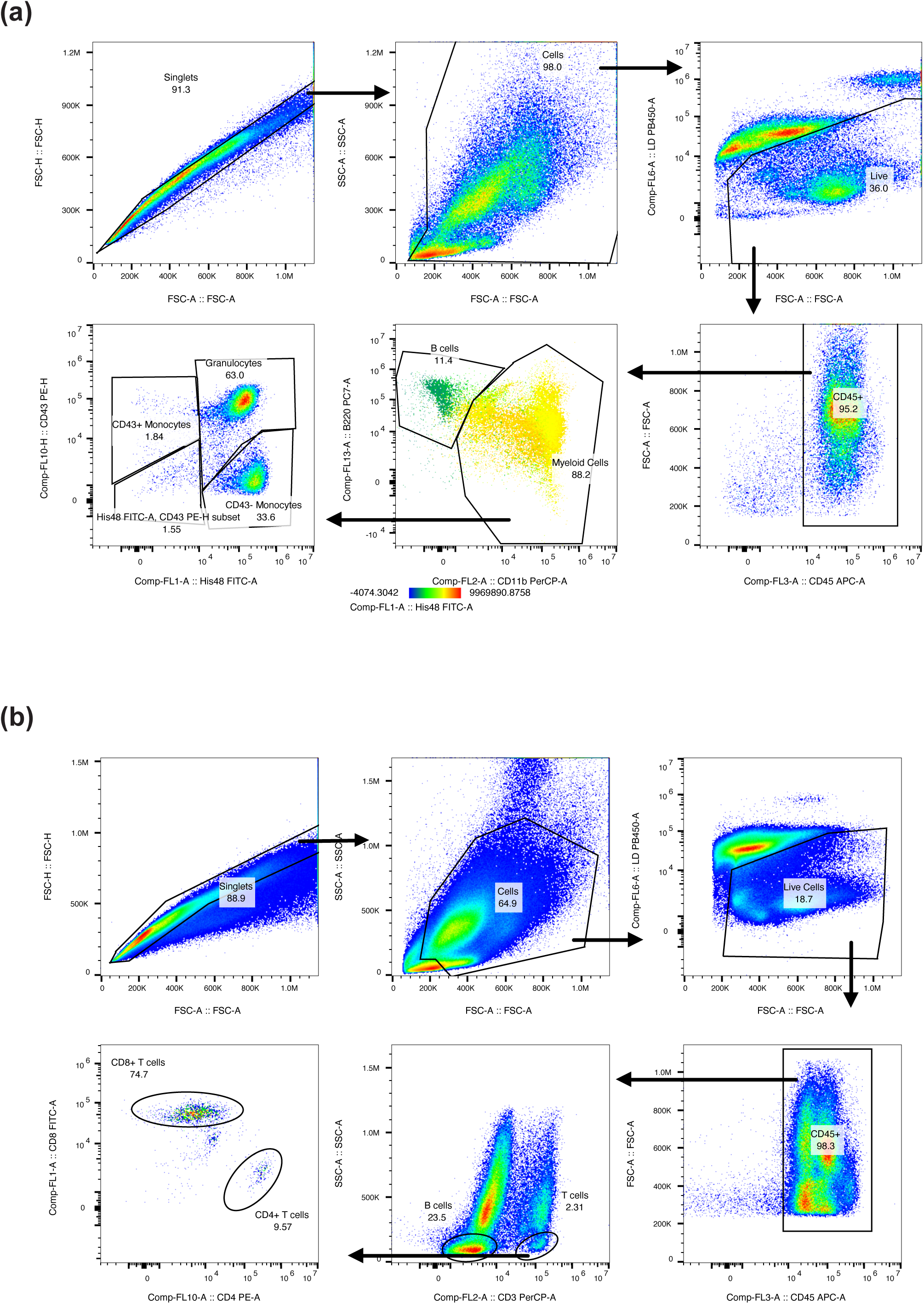
*Polychromatic flow cytometry gating strategy.* Myeloid (A) and lymphoid (B) compartments of bone marrow collected from 12- and 24-week old male and female F1 offspring was examined by flow cytometry.

**Supplementary Figure 2.**
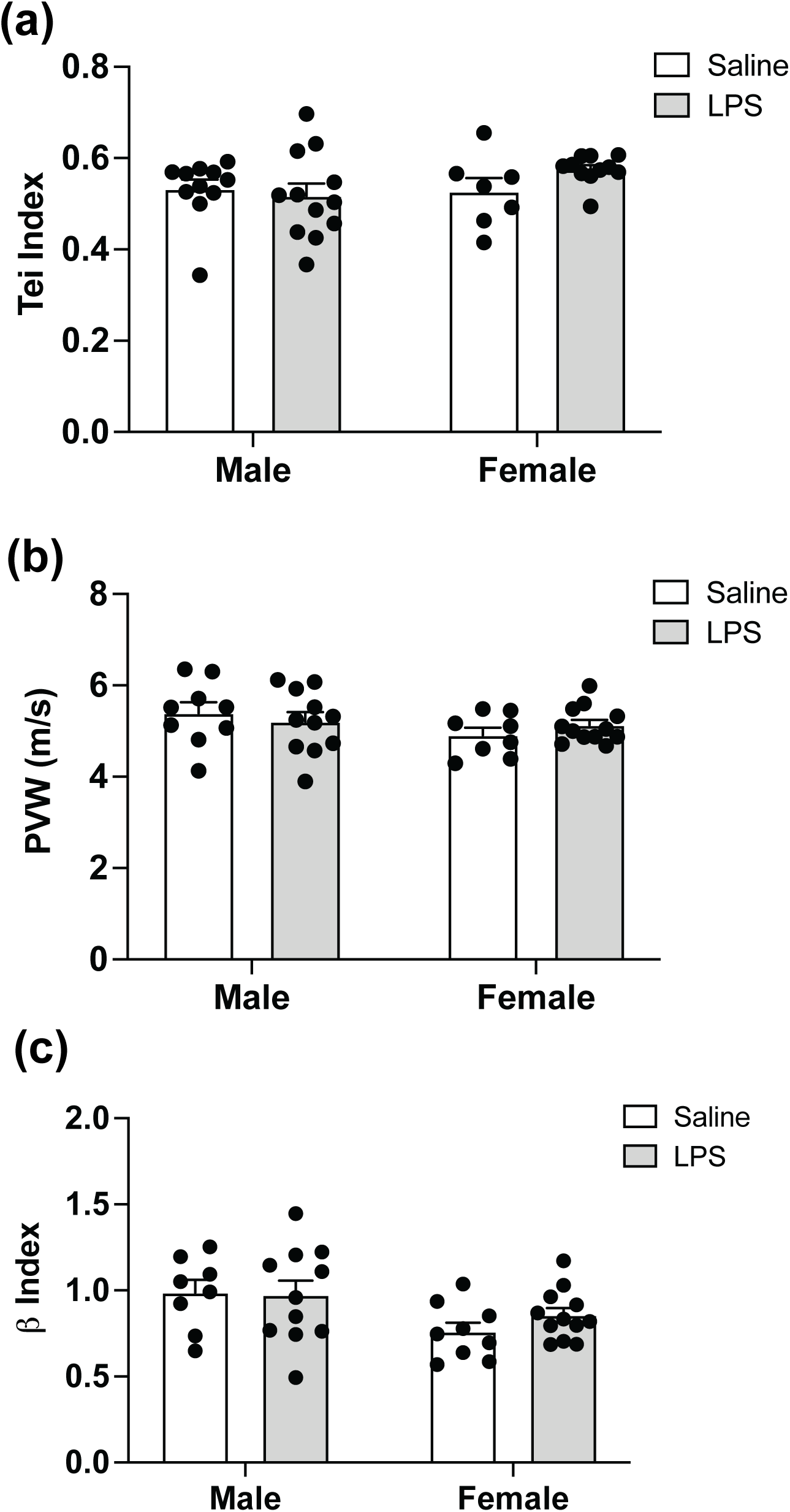
*Effect of aberrant maternal inflammation on F1 cardiac parameters.* Tei index *(a)* was used to evaluate global systolic and diastolic function.B. Pulse-wave velocity (PWV; *b*) was measured to determine aortic wall stiffness. β index normalization *(c)* was calculated to eliminate the influence of diastolic pressure. Each filled circle represents an individual rat.

**Supplementary Figure 3.**
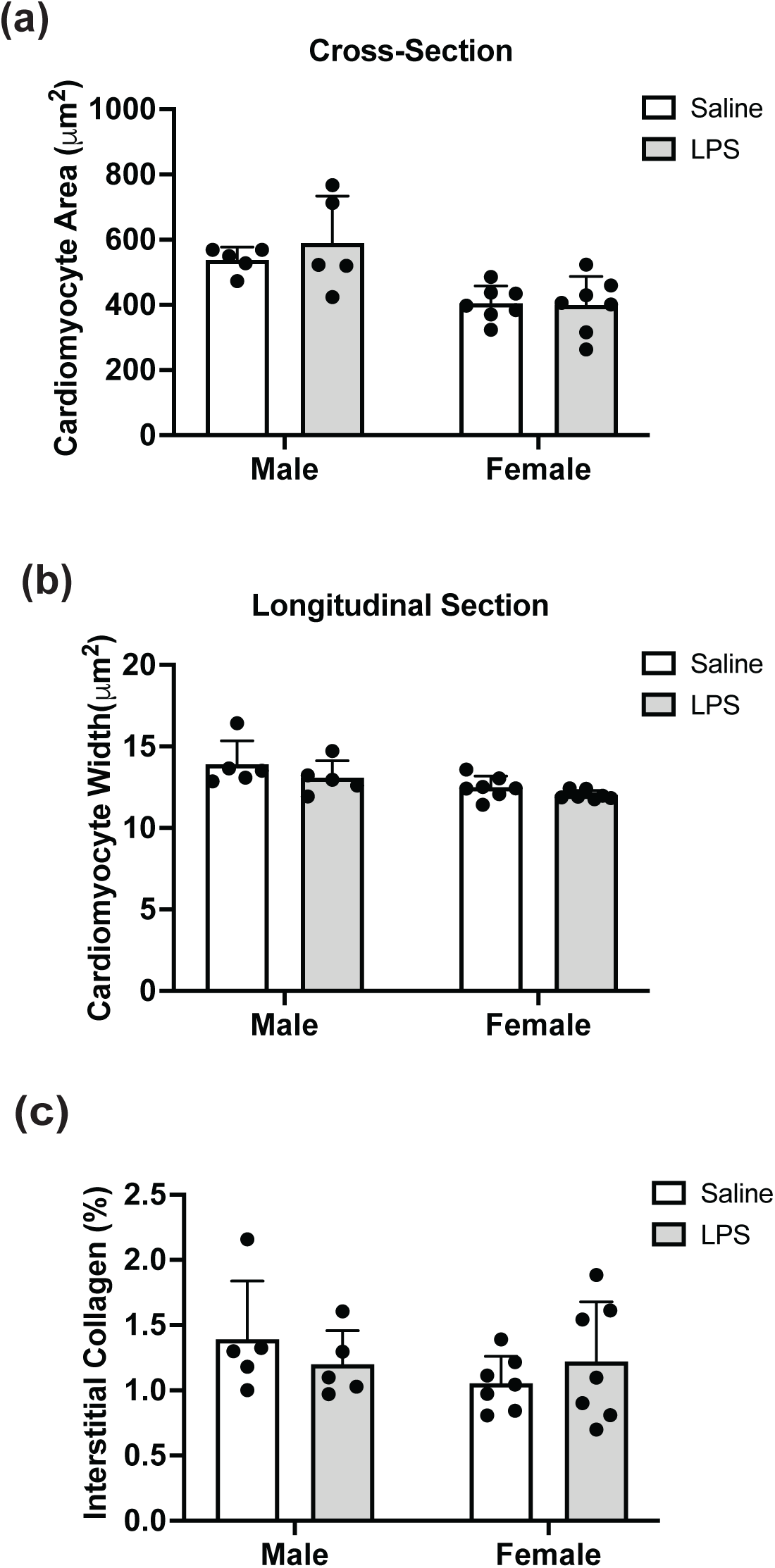
*Effect of aberrant maternal inflammation on offspring cardiac histology.* Cardiac histology including cardiomyocyte area *(a)* and width *(b)*, and interstitial collagen fiber deposition *(c)* was assessed by conventional histological methods at 24 weeks of age. Data are represented as mean ± SEM (n = 5). Each filled circle represents an individual rat.

## Notes

### Competing Interest Statement

The authors have declared no competing interest.

